# A Bayesian decision-making model of implicit motor learning from internal and external errors

**DOI:** 10.1101/2025.01.30.635749

**Authors:** Hyosub E. Kim, Romeo Chua, Davin Hu

## Abstract

A key challenge for the sensorimotor system is deciding which errors to learn from and which to ignore. Recent work has shown that humans are remarkably precise in parsing movement errors into internally- and externally-generated components for this purpose: Participants automatically ignore internally-generated reaching errors caused by motor noise, yet implicitly adapt to size-matched externally-generated errors caused by visuomotor rotations (Ranjan & Smith 2018, 2022). Following replication of these results with 16 neurotypical adults, we formalized our understanding of this behavior with a novel Bayesian decision-making model. The Parsing of Internal and External Causes of Error (PIECE) model frames adaptation as a process of causal inference regarding the source of error, with the magnitude of motor corrections reflecting a combination of state estimation and the observer’s degree-of-belief that their movement was externally perturbed. Thus, PIECE presents a challenge to a class of computational models that frames adaptation as a process of re-aligning the perceived hand position with the movement goal. We show that only PIECE can capture the precise parsing of internal versus external errors observed. Combined, this work provides a normative explanation of how the nervous system discounts intrinsic motor noise and adapts to perturbations, keeping movements finely-calibrated.

## Introduction

Not all motor errors are created equal. Imagine reaching for your morning coffee, but instead of your fingers landing gracefully on the mug, they unintentionally touch down with more force than intended and tip it. This error could be the result of intrinsic motor variability (i.e., noise), the random internally-generated fluctuations in movement parameters that occur even during overlearned movements like reaching. In such a case, it would be counterproductive for your motor system to recalibrate its sensorimotor mapping. Adapting to motor noise, or internally-generated error (IGE), which by definition is unpredictable, would result in highly unstable behavior and potentially amplify the magnitude of future errors even during unperturbed movements (Chaisanguanthum et al., 2014; Faisal et al., 2008; van Beers, 2009; van Beers et al., 2013). On the other hand, if your reach is inaccurate because of an external perturbation, such as the extra inertia from a heavy watch that you just put on, then it is critical to recalibrate your sensorimotor mapping in order to account for the added weight and keep future movements accurate and precise. The sensorimotor system is constantly tasked in this manner with deciding which errors to learn from and which to ignore.

While from a strictly logical viewpoint, it is clear the motor system should not adapt to motor noise but should learn from externally-generated error, what is the empirical evidence that this is actually the case? Perhaps the strongest evidence comes from a remarkable study by Ranjan & Smith which showed that humans are able to accurately parse total movement error into its constituent parts: the error component due to motor noise, or internally-generated error (IGE), and the error component due to an external perturbation, or externally-generated error (EGE). In their study, participants experienced small, randomized visuomotor rotations in which the cursor feedback was rotated relative to their actual hand trajectory by *±*2° or *±*4° on every other trial (Ranjan, 2022; Ranjan and Smith, 2018). Despite the distribution of rotations (EGEs) being matched to baseline motor variability, participants demonstrated robust implicit single-trial adaptation to only the externally-generated component of the total error while effectively ignoring their IGE. That is, immediately following a perturbation trial, the subsequent adaptive response was opposite in direction and proportional to the rotation size (EGE), while also statistically independent of the IGE magnitude. Consistent with this finding, a study of saccadic eye movements also reported more robust adaptation to EGE than IGE (Collins and Wallman, 2012), and we have also shown that visually-clamped errors (Morehead et al., 2017) as small as 1° elicit robust adaptation (Kim et al., 2018). Combined, these studies indicate that, for the motor system, the source of the error determines the implicit adaptive response.

According to standard theories of adaptation, the mismatch between predicted and actual sensory consequences of a motor command, called a “sensory prediction error”, is the main driver of implicit adaptation (Kim et al., 2018, 2019; Mazzoni and Krakauer, 2006; Morehead et al., 2017; Shadmehr et al., 2010; Tseng et al., 2007). The neurophysiological underpinning of sensory predictions is the efference copy, also known as “corollary discharge” (Carriot et al., 2013; Sommer and Wurtz, 2008; Sperry, 1950; Von Holst and Mittelstaedt, 1950), which refers to the copy of the motor command that gets sent from motor cortices to subcortical and sensory regions of the brain. Importantly, computational theories of adaptation typically assume that the sensory prediction is centered on the motor goal, or more specifically, the explicit aiming location, i.e., the path the end effector would travel in the absence of any IGE or adaptation (Mazzoni and Krakauer, 2006; Taylor and Ivry, 2011; Taylor et al., 2014). However, Ranjan & Smith’s study provides compelling behavioral evidence that the sensory prediction also contains information regarding the amount of central motor noise associated with each motor command (Churchland et al., 2006). They argue that the brain generates a highly accurate sensory prediction of the hand’s actual trajectory (minus any peripheral contributions to movement error), rather than the intended trajectory to the target, which explains how the motor system is able to effectively cancel out IGE from the total error. This refined definition of SPE provides a plausible explanation of how the motor system parses total movement error, but what remains to be found is an overarching computational-level theory that explains this behavior.

The aim of the current study was to understand the computational principles underlying such discriminative and finely-tuned adaptive responses. In parallel with replication of the main results of Ranjan and Smith, we formalized our understanding of this behavior by developing a novel Bayesian model, referred to as the Parsing of Internal and External Causes of Error (PIECE) model. Similar to prior Bayesian models of motor adaptation (e.g., Wei and Körding, 2009) and error detection (Gaffin-Cahn et al., 2019), causal inference regarding whether the feedback was perturbed or not is central to the PIECE model. However, PIECE is structurally different from these models in that it assumes that, in addition to vision and proprioception, the observer also has access to an efference copy-based cue of the motor command which includes associated central motor noise (Ranjan and Smith, 2018; Szarka et al., 2024). The PIECE model conceptualizes adaptation as a process of utilizing all three cues to form a posterior belief regarding the presence (or absence) of a perturbation. Critically, the adaptive motor output reflects the estimated perturbation size, which is inferred by the observer by combining their prior on the perturbation with their visual likelihood, weighted by this degree of belief. This framework runs counter to a powerful class of computational models that frames adaptation as a process of aligning the perceived hand position with the movement goal (Tsay et al., 2022; Wei and Körding, 2009; Zhang et al., 2024). The present study challenges such a view, as only the PIECE model could accurately capture the precise parsing of IGE and EGE observed. Our work instead supports a normative view of implicit adaptation in which prediction, perception, and error correction are unified.

## Results

### Differential adaptation to IGE-EGE

We began our study by attempting to replicate the main findings of Ranjan and Smith regarding the sensorimotor system’s decomposition of total error into internally- and externally-generated components (Ranjan and Smith, 2018). We tested 16 healthy, neurotypical young adults on a visuomotor task that involved fast point-to-point reaches to a visual target while controlling a small white cursor, which, depending on the trial, could be rotated by 0°, 2°, or *±*4°. These rotations were pseudorandomized across the experimental block (Fig. 1a shows the experimental set-up) and were always surrounded by null (no rotation) trials. For our analyses, the overall motor error was decomposed into its internally- and externally-generated parts. The internally-generated error (IGE) is defined as the random error that occurs on every reach (i.e., any angular deviation of the hand trajectory from the target), and on unperturbed reaches, serves as the only contribution to total error. IGE is the result of intrinsic motor variability plus any intrinsic bias in reach direction (Wang et al., 2024). Externally-generated errors (EGE) are caused by external perturbations (e.g., visuomotor rotations) and are under experimental control (Fig. 1b). In this experiment, EGEs were defined as the visuomotor rotations, and thus small and statistically independent of IGE. The intention behind utilizing such small perturbations was to keep IGE and EGE on an equal footing in terms of their magnitudes. With each perturbation trial being surrounded by null trials, adaptation was quantified as the difference in reach angle between the trial immediately following versus immediately preceding the perturbation trial: adaptation_*t*_ = hand_*t*+1_ *−* hand_*t−*1_, where *t* indexes the perturbation trial. Combined, these methods allowed us to easily dissociate the influence of IGE versus EGE on implicit adaptation on a trial-by-trial basis (see Data Analysis).

**Figure 1:**
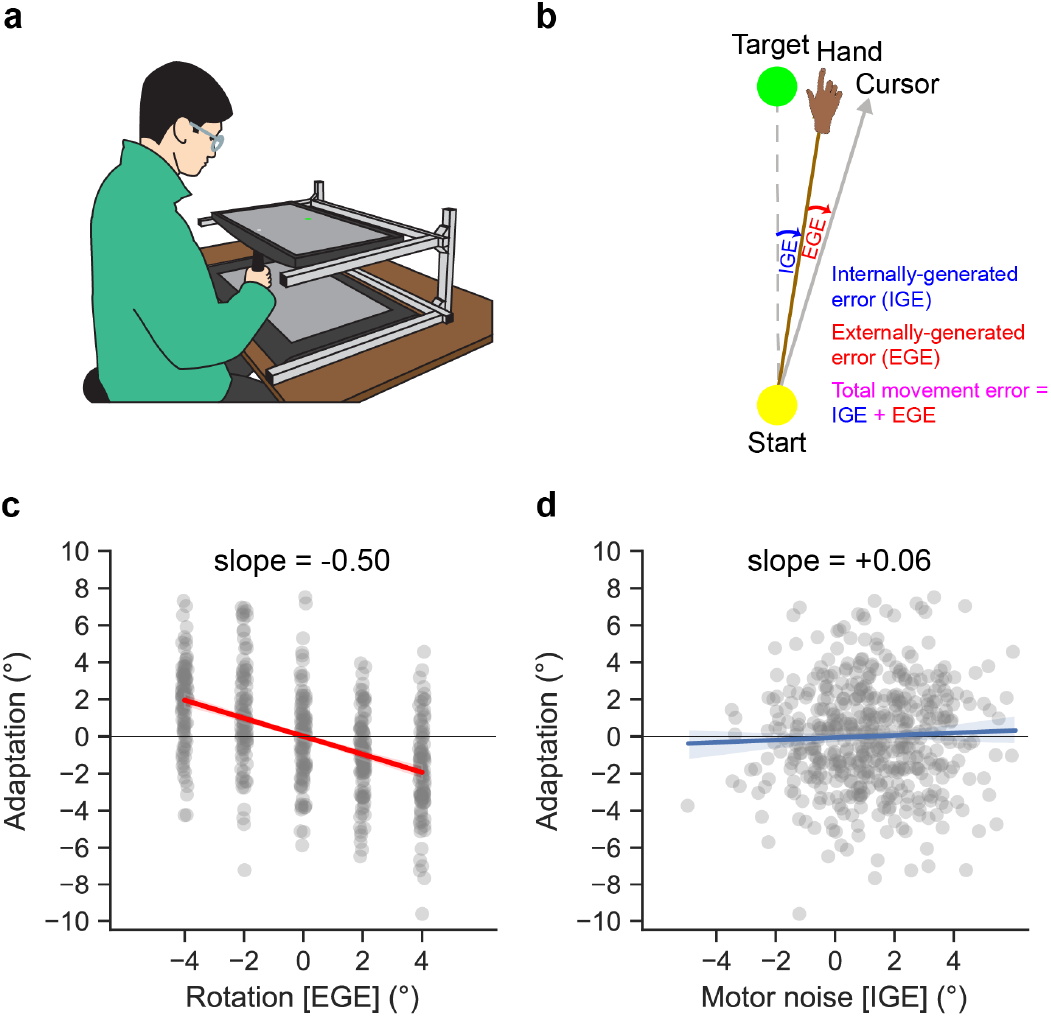
(a) Basic experimental set-up involved participants making quick point-to-point reaches by sliding a stylus along a graphics tablet. Participants could not see their hand, as experiments were conducted in a darkened room and the horizontally-oriented monitor blocked vision of their hand. (b) Schematic showing how we operationally defined the two sub-components of the total error. The internally-generated error (IGE) is equivalent to motor noise, or how far off the intended aim (i.e., the target) the reach was. The externally-generated component (EGE) refers to the external perturbation—in this case, the magnitude of the visuomotor rotation. (c) Individual participant data showing adaptation as a function of EGE (gray dots represent single-trial adaptive responses). There is a clear, distinct response to EGE. (d) When the same adaptive responses are plotted as a function of IGE, there is no discernible relationship between the variables. Note that the data are shifted slightly rightwards due to a small counter-clockwise reaching bias. The shaded regions in (c) and (d) represent bootstrapped 95% CIs (difficult to see due to low variance).

In Fig. 1c-d we present data from an example participant. Separately plotting adaptation to EGE and IGE shows a clear dicohotomy in the responses to these two different types of error. Consistent with the findings of Ranjan & Smith, there is robust adaptation to the externally imposed visuomotor rotations but no evidence of adaptation to spontaneously generated errors due to motor noise, which spanned a similar range as the EGEs. This dissociation between adaptive responses to EGE versus IGE was consistent across our entire sample.

We performed a similar set of group-level analyses as Ranjan & Smith to better understand the population-averaged adaptive responses to errors. Briefly, we first binned the data based on the level of EGE, and then for each of the five levels of EGE, we further binned the data based on the level of IGE into quintiles (see Methods). This procedure was applied separately for each participant. The results of these analyses are presented in Fig. 2a-b, where each point represents the average across every participant’s mean response for the corresponding EGE/IGE level. As seen in Fig. 2a, there was a clear linear response to EGE (black dashed line, mean slope=-0.594, bootstrapped 95% CI: [−0.643, −0.545], *r*^2^=0.991, *p <* 10^*−*24^), with over 99% of the variance in the EGE/IGE grid being explained by the level of EGE. In stark contrast, Fig. 2b shows that there was no systematic relationship between adaptation and IGE, with less than 0.2% of the variance in the EGE/IGE grid being explained by IGE (dashed black line, mean slope=-0.043 [-0.071, 0.000], *r*^2^=0.002, p=0.82).

**Figure 2:**
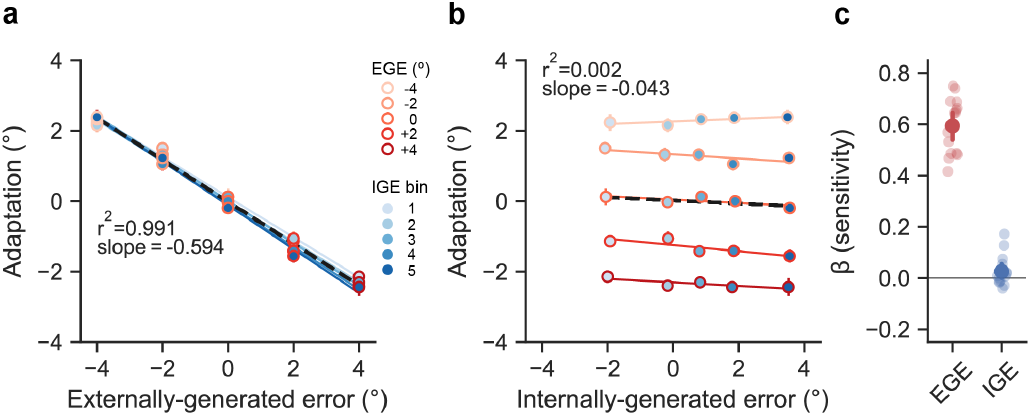
(a) Population-averaged adaptive responses were binned based on the level of EGE (shade of red) and IGE (shade of blue). The filled-in circles represent the mean of each quintile and are plotted as a function of EGE in (a), and as a function of IGE in (b) (data are shifted right due to a small counter-clockwise reaching bias of less than 1° across participants). The vast majority of the variance in adaptive responses (99.1%) was explained by EGE. (c) Linear regression coefficients from an analysis of unbinned data (signs are flipped for ease of comparison). Error bars represent bootstrapped 95% CIs and are quite small in (a) and (b). More translucent dots in (c) represent individual participants.

To further quantify the impact of EGE versus IGE on implicit adaptation, we performed a bivariate regression analysis of the unbinned data from each participant, using EGE and IGE as predictor variables. The coefficients of each predictor (sign-flipped for more convenient comparisons) were markedly different (mean difference = 0.577, 95% CI for difference scores: [0.508, 0.645]; *t*_15_ = 17.9, *p <* 10^*−*10^), reflecting the high sensitivity of each participant to EGE (0.599[0.550, 0.648], *t*_15_ = 23.1, *p <* 10^*−*12^), with correspondingly low sensitivity to IGE (0.022[*−*0.004, 0.052]; *t*_15_ = 1.51, *p* = 0.153). In line with the original results of Ranjan & Smith, our behavioral results highlight the motor system’s clear parsing of total movement error into internally- and externally-generated components, effectively discounting the former and learning from the latter.

### The Parsing of Internal and External Causes of Error (PIECE) Model

The primary purpose of the current study was to understand the computational principles underlying the motor system’s incredibly accurate parsing of movement error into IGE and EGE. Towards this aim, we have developed a Bayesian model that optimally combines all available sensory cues in order to form a posterior belief regarding the presence (or absence) of a perturbation, with motor output reflecting the weighting of the estimated perturbation size by this degree of belief. As seen in the graphical generative model of Fig. 3, the observer has access to cues regarding hand position from vision (*x*_*v*_), proprioception (*x*_*p*_), and an efference copy-based sensory prediction of hand position (*x*_*u*_), which we refer to as a “motor prediction” since it stems from the motor command. The proprioceptive and motor prediction cues are unbiased estimates of the actual hand position, *x*_*h*_. That is, 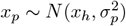, and 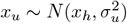. During perturbed trials with a visuomotor rotation, the visual cue is offset from the actual hand location by the size of the visuomotor rotation, 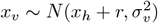, while, by definition, *r* = 0 during unperturbed trials. Based on a recent adaptation study that showed that participants’ visual uncertainty increases as a function of the error size, we also assume the observer’s visual uncertainty, *σ*_*v*_, increases as a linear function of the distance of the feedback cursor from the target (i.e., the target error, *e*): *σ*_*v*,*t*_ = *α* + *β · e*_*t*_, where *t* indexes the trial number (Zhang et al., 2024).

**Figure 3:**
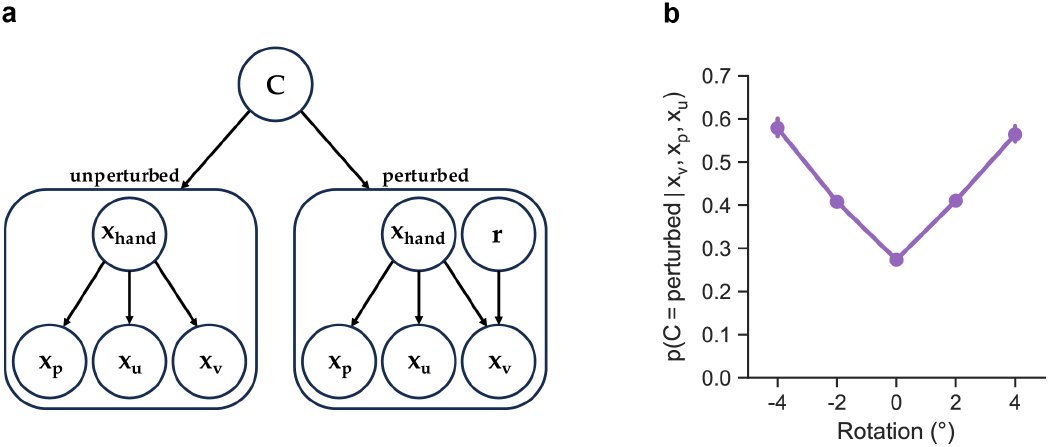
(a) Generative model for PIECE. In this model, the observer uses proprioceptive (*x*_*p*_), motor predictive (*x*_*u*_), and visual cues (*x*_*v*_) to compute posterior probabilities of *C*, the causal node, which determines whether the feedback was perturbed or not. In the unperturbed case visual feedback is a function of the actual hand position only, whereas in the perturbed case visual feedback is a function of both the actual hand position, *x*_hand_, and the rotation, *r*. On a trial-by-trial basis, the posterior over *C* weights the observer’s posterior estimate of *r*, which is computed by combining the prior on the rotation and the likelihood associated with the visual measurement. (b) The posterior, *p*(*C* = pert|*x*_*v*_, *x*_*p*_, *x*_*u*_), as a function of the rotation size (EGE) for the same participant shown in Fig. 1. This function was derived by first finding the maximum-likelihood estimates (MLEs) of this participant’s data with PIECE (i.e., the parameter set that maximizes the probability of their observed data), and then using these MLEs to simulate behavior across the experimental protocol and recording the posterior estimates.

With respect to the other nodes in the generative model, the observer’s prior on hand position, *x*_*h*_, is normally distributed: 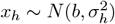, where *b* represents any directional bias in reach angle relative to the target (Wang et al., 2024; see Fig. 2b) and *σ*_*h*_ is equal to the participant’s intrinsic motor variability. This prior on *x*_*h*_ reflects the fact that human participants reach in a manner consistent with internal knowledge of the distribution of their intrinsic motor noise (Trommershauser et al., 2003; Trommershäuser et al., 2008). The observer also has a prior for the rotation magnitude, 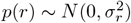. We assume the observer believes the distribution of perturbations to be normally distributed, with *σ*_*r*_ representing the range of perturbation sizes they believe is plausible. The prior used by the observer on the causal node, which reflects their degree of belief in whether the reach was perturbed or not, matches the statistics of the task and is flat: *p*(*C* = perturbed) = *p*(*C* = unperturbed) = 0.5.

For our model-fitting, we calculated each participant’s motor variability during the last 50 trials of baseline reaches and used this value for *σ*_*h*_. To further limit the number of free parameters, rather than fitting *α* and *β* of the visual uncertainty function, we used the reported mean values from the study by Zhang and colleagues (see Table S1 from Zhang et al., 2024). In total, PIECE has 3 free parameters: *σ*_*r*_, *b* and *σ*_combined_, the latter representing the standard deviation that results from combining random variables *x*_*p*_ and *x*_*u*_. Briefly, while the observer may have access to separate proprioceptive and motor prediction cues, from the experimentalist’s perspective, *x*_*p*_ and *x*_*u*_ are not dissociable due to their both being internal measurements centered on the true hand location. See Modeling Analysis for the derivation of *σ*_combined_ from *x*_*p*_ and *x*_*u*_.

#### Causal inference in PIECE

Beginning with Bayes’ Rule, the observer combines the prior, *p*(*C*), and the likelihood, *p*(*x*_*v*_, *x*_*p*_, *x*_*u*_|*C*), to compute the posterior for the causal node, *C*:

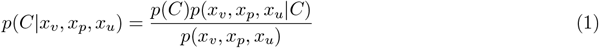

As in most Bayesian cue combination models, we assume the cues are conditionally independent and their respective Gaussian distributions (likelihoods) are multiplied. Performing causal inference and computing the posterior for the unperturbed case (abbreviated as “*¬*pert” below) requires marginalization over the hand position, *x*_*h*_:

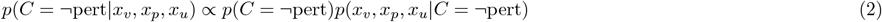

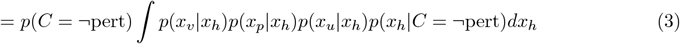

The posterior for the perturbed case (abbreviated as “pert”) requires marginalization over the hand position, *x*_*h*_, and the rotation magnitude, *r*:

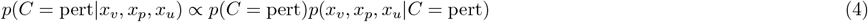

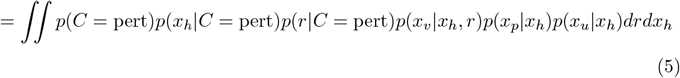

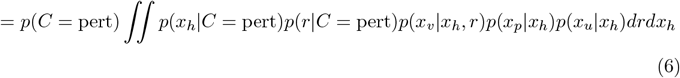

The denominator in Eqn. 1 (i.e., the “marginal likelihood”) serves as a normalization factor that is common to both *C* = *¬*pert and *C* = pert and thus does not need to be formally computed. Instead, as a final step, we can divide the unnormalized posteriors for unperturbed and perturbed world states by their sum to normalize them.

The posterior on *C* defines the observer’s relative degrees of belief in both the unperturbed and perturbed world states. Fig. 3b shows the posterior on *C* = pert as a function of the rotation size (EGE) for the same participant shown in Fig. 1c-d. As seen in this figure, the observer assigns more credibility to the hypothesis that their reach was perturbed as EGE increases. However, some degree of belief gets assigned to the perturbed hypothesis even during unperturbed trials, highlighting the probabilistic nature of motor adaptation (Berniker and Kording, 2008). In the PIECE model, these degrees of belief are combined with optimal state estimation of the perturbation magnitude.

#### State estimation in PIECE

The observer updates their state estimate of the perturbation with each observation (measurement). This *estimate* is denoted 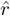, whereas *r* represents the true perturbation, which is unknown to the observer. For this state estimation step, we assume the observation on a given trial, *z*_*t*_, which is distributed as 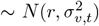, is an unbiased measurement of the perturbation on that trial (i.e., the actual rotation size; Burge et al., 2008; Haith and Krakauer, 2013; Wei and Körding, 2010), as opposed to *x*_*v*,*t*_, which is a function of the perturbation and the true hand location. Since the perturbations varied randomly in this experiment, we assumed the trial-by-trial effects were independent of each other and that there was no memory across trials (Ranjan, 2022; Zhang et al., 2024).

To update their estimate of the rotation size, the observer should combine their measurement with their prior on the rotation. Doing so leads to the proportion of the error that is corrected for on each trial being specified by *K*_*t*_ (Eqn. 10), where 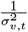 and 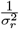 represent the measurement and prior precisions, respectively (Burge et al., 2008; Haith and Krakauer, 2013; Korenberg and Ghahramani, 2002). The key intuition here is that the learning rate (*K*_*t*_) should decrease with increasing visual uncertainty (*σ*_*v*,*t*_), and conversely, increase the more volatile the observer believes the environment is, represented by a higher value of *σ*_*r*_.

Taken together, the overall state estimate of the perturbation on trial *t*, 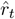, is a linear combination of the state estimates under the unperturbed and perturbed world state hypotheses:

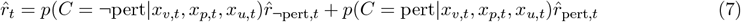

In the unperturbed world state, there is no perturbation and so *r* = 0 by definition. Thus, the contribution of 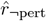 to the overall state estimate is zero. The full expression for how much the observer adapts their overall state estimate, 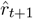, is therefore expressed as follows:

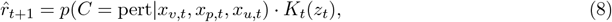

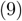

where

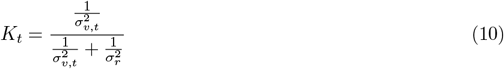

Lastly, we assume the observer’s motor output is a reflection of their estimate of the perturbation (thus opposite-signed) plus their intrinsic motor bias and motor noise:

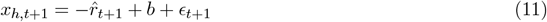

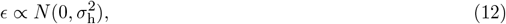

where *σ*_*h*_ is measured during baseline reaches.

As expressed in Eqn.11, the observer’s motor output reflects the combination of the estimated rotation size and the strength of their belief in the presence of an external perturbation. Thus, by combining Bayesian cue combination, causal inference, and state estimation, the PIECE model unifies competing notions of the computational goal of implicit adaptation.

### Model-based analyses

We evaluated the performance of four computational models of adaptation on our behavioral data: PIECE, the Proprioceptive Recalibration Model (PReMo) (Tsay et al., 2022), the Perceptual Error Adaptation Model (PEA) (Zhang et al., 2024), and the Relevance Estimation Model (REM) (Wei and Körding, 2009). The models competing with PIECE, collectively referred to here as “hand-to-target alignment” models, were chosen because of their previously demonstrated ability to explain a wide range of adaptation phenomena, the fact that they are among the most prominent Bayesian, or Bayesian-inspired, models of adaptation, and, importantly, because they each frame the computational goal of adaptation as one of aligning the perceived hand position with the movement goal. While the development of the PIECE model was informed by these other models, in stark contrast to them, PIECE posits the computational goal of adaptation as being one of correcting for the externally-generated error (EGE). By objectively comparing the performance of these four models, we can adjudicate between the hand-to-target versus error correction views of adaptation.

The high-level intuition of PReMo is that mismatches between visual, proprioceptive, and efference copy-based cues result in a recalibrated estimate of the “felt” hand position, and implicit adaptation serves to minimize any misalignment between this perceived hand position and the motor goal, or target. PEA also posits adaptation as being driven by a perceptual error. The derivation of the perceived hand position in PEA follows optimal integration principles (Ernst and Banks, 2002; Landy et al., 1995), and as mentioned earlier, the model assumes visual uncertainty increases with error size. REM is closest in structure to PIECE, requiring a causal inference step. However, similar to PReMo and PEA, REM also assumes realignment of the estimated hand position with the target as the primary goal of adaptation.

We started our analyses by using maximum likelihood estimation to separately fit each individual’s data with all four models. After finding the set of parameters that maximized the probability of each participant’s data, we computed and compared Bayesian Information Criterion (BIC) scores (Schwarz, 1978). Fig. 4 shows the results of this analysis, with panel A showing ΔBIC scores for each model when benchmarked to PReMo’s BIC scores for each individual. (We chose PReMo as the primary comparison model based on our involvement in its development and also because it is most explicit in promoting the hand-to-target alignment idea.) Clearly, PIECE had the lowest (best) BIC scores, indicating that out of all of the models it most successfully captured the parsing of total movement error into IGE and EGE demonstrated by all participants. When directly comparing BIC scores between PIECE and each of the other three models, one can see that PIECE was not just the best overall model, but that it unanimously did the best job of fitting each individual’s data.

**Figure 4:**
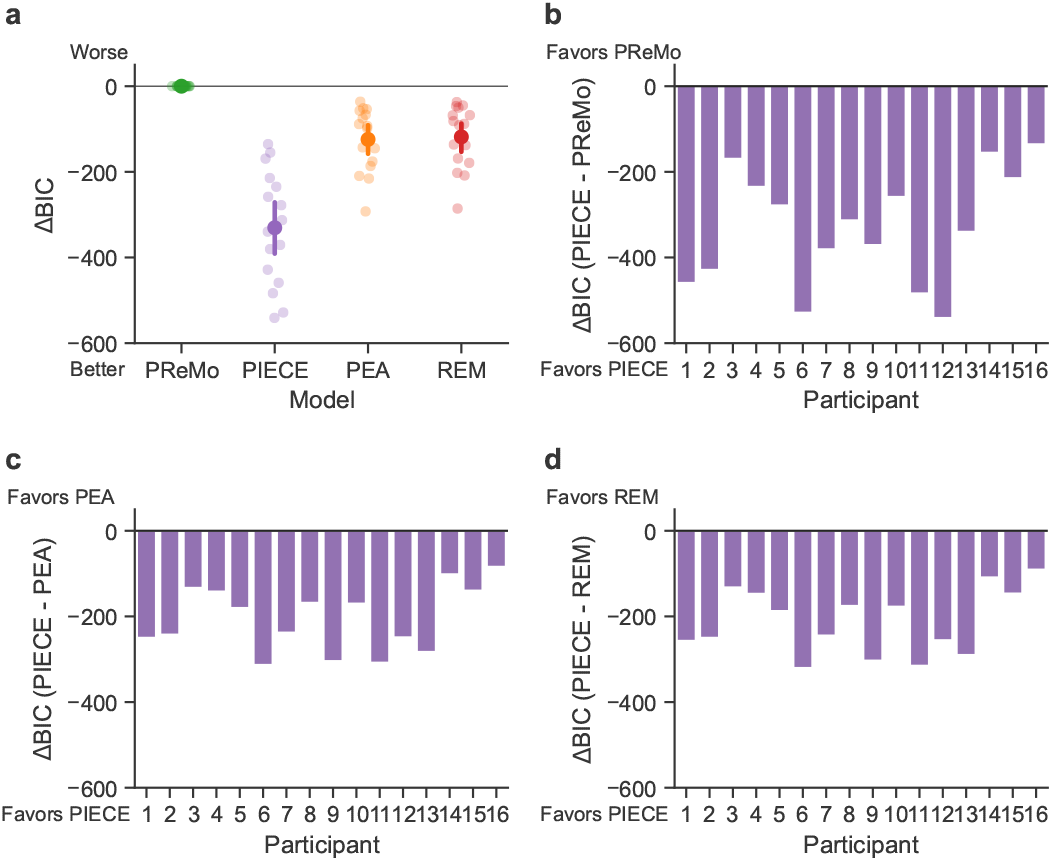
(a) Direct comparison of BIC scores of each model to PReMo’s BICs. Lower scores indicate a better model fit. (b-d) For all 16 participants, PIECE outperformed each of the other three models.

In addition to objective model selection criteria, we also performed a posterior predictive check of our models by generating simulated data with the best-fit parameters (see for parameter values). The logic of this procedure is that any successful model should be able to approximate the observed data (Wilson and Collins, 2019). Fig. 5 shows data from a representative participant, along with simulated data using each model’s maximum likelihood estimates of parameter values for this participant. Only the PIECE model successfully captured the sensorimotor system’s ability to robustly adapt to small external perturbations while simultaneously discounting motor noise. Combined, we see that PIECE is the most appropriate model for these data based on objective model selection criteria, and that, qualitatively, it was the only model that could capture participants’ highly accurate decomposition of total error into constituent parts.

**Figure 5:**
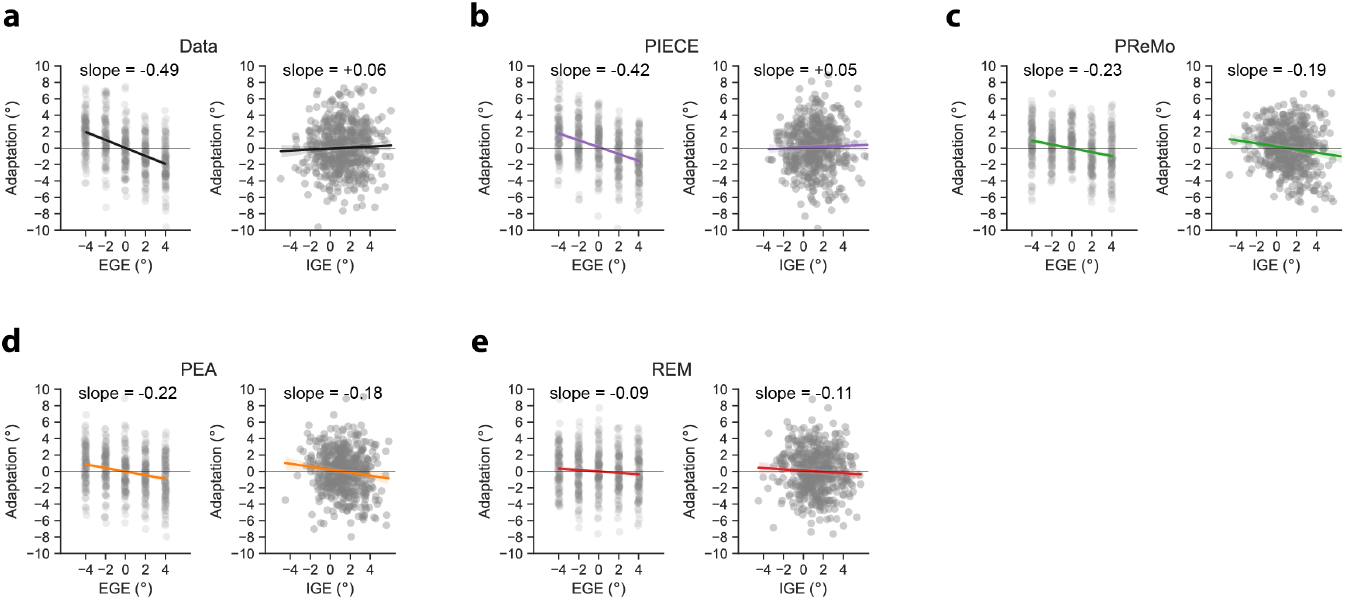
Posterior predictive check via simulation of entire experiment using MLEs of each model’s parameter set. (a) Representative data from a single participant (P13). Adaptive responses are a linear function of EGE and remain statistically independent of IGE. (b) PIECE model mimics the empirical data: Slope values indicate high senstivity to EGE and insensitivity to IGE. (c-e) PReMo, PEA, and REM all fail to capture the accurate parsing of errors into EGE and IGE. Gray dots represent adaptation measures of individual trials. Thick colored lines and shading represent line of best fit and associated bootstrapped 95% CI, respectively. Posterior predictive checks of other 15 participants followed qualitatively similar pattern.

The behavioral data from the current study and from Ranjan & Smith make clear that responses to IGE and EGE are statistically independent. For this reason, any computational framework that posits the goal of adaptation as one of aligning the hand with the target—as PReMo, PEA, and REM do—is bound to fail. While those models capture adaptation to EGE, they also incorrectly predict adaptation to be driven by IGE, as clearly illustrated in Fig. 5. Even on unperturbed trials, the misalignment of the hand with the target due to intrinsic motor variability will trigger an adaptive response in all models other than PIECE. However, a plethora of data now show that human participants do not treat IGE and EGE equally, which only the PIECE model was able to explain.

## Discussion

As first reported by Ranjan & Smith, the adaptation system shows a remarkable capability to filter out IGE from total error in order to implicitly recalibrate its sensorimotor mapping in response to external perturbations (Ranjan, 2022; Ranjan and Smith, 2018). Our successful replication of these results supports this finding and points to its robustness. To provide a computational-level explanation of the observed error parsing behavior (Blohm et al., 2020; Krakauer et al., 2017; Marr, 2010), we developed the PIECE model. PIECE is a Bayesian decision-making model that incorporates ideas of cue combination and causal inference, and takes inspiration from the three models it was compared to in this study, PReMo, PEA, and REM. In contrast to these other models, though, PIECE returns to classical ideas of adaptation as a process of state estimation with regard to the external perturbation (Krakauer et al., 2019; Shadmehr et al., 2010). Indeed, if one were to simply cast adaptation as a process of proportionally updating motor output by a fraction of the experienced rotation size, as in standard state-space models of adaptation (Donchin et al., 2003; Smith et al., 2006; Thoroughman and Shadmehr, 2000), we could recapitulate the differential adaptation to IGE-EGE observed in our study. However, such a model does not tell us *why* or *how* the sensorimotor system decides which errors to adapt to, and which errors to ignore, not to mention other limitations of state-space models (Kim et al., 2018; Krakauer et al., 2019). The unique advance offered by the PIECE model is that it provides a coherent computational-level explanation of how the sensorimotor system parses motor errors into their respective internally- and externally-generated components in order to ignore the former and adapt to the latter.

### Implicit adaptation is not driven by perceptual error

PReMo and PEA are well-developed models that explicitly formulate the computational goal of adaptation as one of realigning the perceived hand position with the target. Both models posit that in a perturbed environment, such as a visuomotor rotation, the mismatch between sensory and motor prediction-based cues results in a proprioceptive shift towards the visual cursor, generating the driving signal for adaptation (Tsay et al., 2022; Zhang et al., 2024). Although these models differ in their explanation of how this shift occurs and what constraints are placed on it, both are incapable of accommodating the error parsing observed here and first reported by Ranjan & Smith (Ranjan and Smith, 2018). As our data and analyses make demonstrably clear, the perceived hand position, and its relationship to the target, cannot serve as the primary driver of single-trial adaptation. Fig. 5 shows that, while these models can accommodate participants’ responses to EGE, they incorrectly predict adaptation in response to IGE, since the hand is by definition misaligned with the target when there is motor noise, even in the absence of a perturbation. These models fall short, in part, because they do not incorporate causal inference.

### The role of causal inference in adaptation

Our results indicate that the implicit adaptation system performs causal inference with respect to the sources of the observed total error. Bayesian causal inference models were originally developed to explain why auditory and visual cues are sometimes fused, forming a coherent percept, and why they are sometimes perceived as having two separate sources (Körding et al., 2007; Sato et al., 2007). In the realm of motor adaptation, causal inference was formalized by Wei & Kording with their Relevance Estimation Model (REM)(Wei and Körding, 2009). Although REM and PIECE are structurally similar in terms of causal inference, the posterior on whether the reach was perturbed or not is used in nearly opposite ways in the two models—to correct for errors that are believed to be self-generated (i.e., IGE), in the case of REM, and to correct for errors that are believed to be externally-generated (i.e., EGE), in the case of the PIECE model.

According to REM, visual errors that fall within the range one should expect to observe, based on normal motor variability, are deemed more relevant and thus require an adaptive response. In their paper, the authors tested a much larger range of error sizes (≈ 4° − 58°) and were not at all concerned with explaining parsing of total movement error into IGE and EGE. And similar to nearly all adaptation studies, the methods they used did not allow them to specifically examine responses to IGE (see Methods), so perhaps it should not be surprising that their proposed explanation of adaptation differs from ours. Regardless of these methodological differences, the current data and the PIECE model make clear that, depending on their inferred source, size-matched errors are treated drastically differently by the motor system, and only the external component of error should be corrected for.

### The role of prediction in adaptation

REM includes only proprioceptive and visual cues regarding actual hand position, while both PReMo and PEA also include an efference copy-based predictive cue of hand position. However, in PReMo and PEA the motor prediction is assumed to be centered on the motor goal (i.e., the target), rather than where the central motor system actually directs the hand, which would equal the participant’s explicit aim plus any associated IGE (and, possibly, any implicit adaptation). In this manner, the predictive cues in PEA and PReMo are akin to the prior on hand position in the PIECE model, *p*(*x*_*h*_), which is equal to the actual distribution of unperturbed baseline reaches. In other words, the observer in PIECE possesses knowledge of their own intrinsic motor variability (and bias), an assumption made based on findings from a series of reaching studies examining rapid value-based decision making (Trommershauser et al., 2003; Trommershäuser et al., 2008). In the PIECE model, the combination of the prior on hand position and the predictive cue centered on the actual hand position contribute to effective error parsing.

Based on our model fitting results, the motor prediction cue provides highly accurate and precise information regarding the actual motor command, including noise. The distribution of *σ*_combined_ values across participants was quite tight, ranging from 0.2 to 1.4. This suggests that for many participants, there was very little uncertainty regarding where the hand was actually sent. These values must be interpreted with caution, however, as they actually represent the uncertainty associated with the combination of proprioceptive and motor prediction-based cues, as the two could not be dissociated in this paradigm (see Methods). Regardless, these parameter estimates still suggest a highly accurate and precise prediction, as proprioception is known to be quite noisy. This finding further supports the idea first presented by Ranjan & Smith that, in order to effectively parse errors as small as those used in our paradigms, the precision of the motor prediction must be much greater than that of the movement itself (Ranjan, 2022).

### PIECE and other sensorimotor learning phenomena

PIECE incorporates a key insight from the PEA model related to goal-directed reaching—that visual uncertainty increases with error size. Zhang and colleagues empirically showed that, during reaching, the distance of the cursor from the target will increase visual uncertainty (decrease precision). An important caveat is that the authors were applying this principle to data involving implicit adaptation to visual error clamps, in which participants are explicitly instructed to ignore the cursor feedback and to fixate on the target, and the cursor trajectory is clamped to a specified angle relative to the target. Despite the difference between our tasks, our use of the same linear function for visual uncertainty is justified based on prior work which showed that the eyes automatically direct themselves towards a reaching target (Neggers and Bekkering, 2002). The success of our models further extends the findings of Zhang and colleauges, advancing the notion that whether the cursor feedback is contingent on hand position (as in our study) or not (as in a clamp study), the visual estimate of the cursor position will increase in uncertainty as a function of cursor-to-target distance.

In the Zhang et al study, this increasing visual uncertainty is used to explain non-linear adaptive responses to increasing perturbation sizes. Several studies have now shown that sensitivity to external perturbations is only constant over a narrow range of error size, and that adaptive responses quickly saturate as perturbation size increases (Hayashi et al., 2020; Kim et al., 2018; Marko et al., 2012; Morehead et al., 2017; Wei and Körding, 2009). Importantly, the PIECE model can also account for the saturation phenomenon. As the learning rate (*K*_*t*_) in PIECE is derived from optimally combining prior and likelihood, it is a function of the observer’s prior uncertainty regarding the rotation magnitude, *σ*_*r*_, and their visual uncertainty, *σ*_*v*,*t*_. As visual uncertainty increases with rotation size, the learning rate decreases, helping to capture the saturation of adaptive responses in a principled manner, avoiding the use of an arbitrary free parameter for the learning rate as used in most state-space models. Indeed, when visual uncertainty is combined with the observer’s prior over possible perturbation sizes, PIECE can flexibly accommodate any non-monotonicities in adaptation as a function of error size, as shown in Fig. 6. It should also be noted that if one were to simply add a retention parameter to PIECE (or equivalently, assume temporal dynamics in the state estimation step—see 7), this model could also accommodate the saturation of asymptotic adaptation in response to consistent perturbations (Bond and Taylor, 2015; Morehead et al., 2017; Zhang et al., 2024).

**Figure 6:**
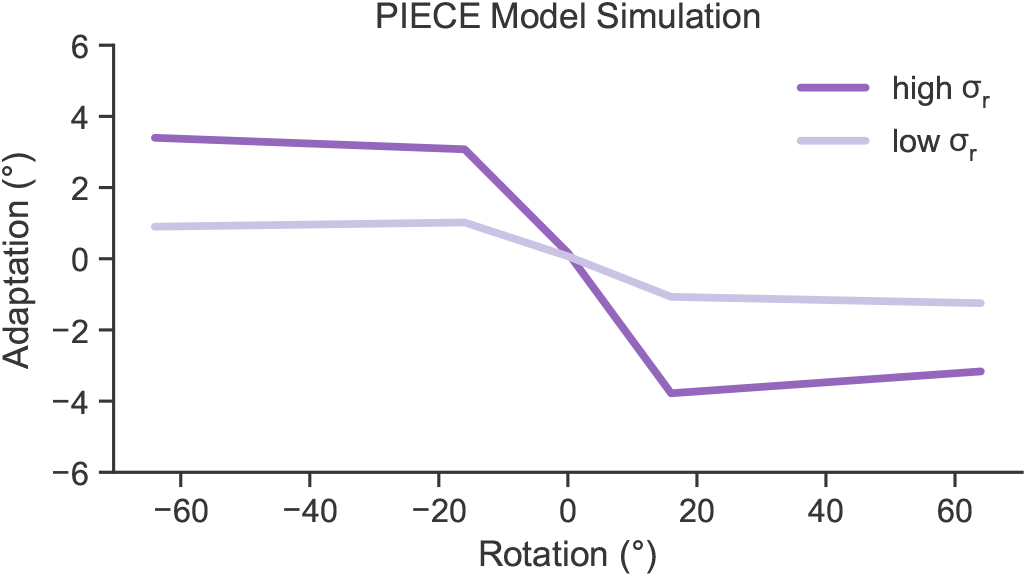
PIECE can accurately capture the non-linear adaptive response to a wide range of error sizes. Here, we simulated responses to *±*4°, *±*16°, and *±*64° errors, as in Experiment 4 from Zhang et al., 2024. Note that both the inflection point and magnitude of adaptation can be flexibly accommodated by PIECE. They are both functions of the observer’s degree of belief that they are perturbed, and therefore depend on *σ*_*r*_ and *σ*_*v*_, the width of the observer’s prior on rotation size and visual uncertainty, which increases with error size (Zhang et al 2024).

## Conclusions

There are three important insights provided by the PIECE model. The first insight is that causal inference is critical for adaptation, both to infer the source of error and also to regulate how much weight to apply to the observer’s estimate of the perturbation, which in turn dictates the magnitude of single-trial adaptation. Without causal inference, other models such as PReMo and PEA are unable to capture the statistical independence between responses to IGE and EGE, since the perceptual error driving adaptation in these models is a function of hand position (IGE). Relatedly, the second insight is that accurate error parsing can only be understood through the lens of state estimation with respect to the EGE, i.e., the perturbation. Despite incorporating causal inference, the REM model also failed to account for the data because it too frames adaptation as a process of inferring the hand position for purposes of aligning it with the motor goal. However, our data and analyses make clear that adaptation responds almost exclusively to EGE. Any minimal response to IGE is attributable to the probabilistic nature of inferring and estimating the source of error—in other words, the motor system is not perfect in its error parsing. Despite this latter point, the third insight is that the internal prediction of where the hand is sent during a reach is highly accurate and precise. This was made clear in the original work of Ranjan & Smith, while PIECE embeds this phenomenon within a Bayesian framework. When considered within the context of high-level motor skills where the margin of error is miniscule, like playing the violin or grabbing our morning coffee in a rush, perhaps it should not be surprising that our sensorimotor system has evolved to keep accurate tabs on where it sends our limbs so that it can finely recalibrate its mapping in response to external perturbations.

## Methods

### Participants

Healthy, young adults were recruited from the University of British Columbia community (N = 16, 8 females; average age = 22.3 years old, range: 19-29). All participants were naive to the purpose of the experiment and provided their written informed consent (consistent with the Helsinki Declaration). Participants were remunerated $15 for their participation. Experimental procedures were approved by the behavioral research ethics board at the University of British Columbia under study ID H23-02324.

### Experimental Set-Up

Participants sat in front of a testing set-up comprised of a horizontally-oriented, 144 hertz refresh rate monitor (53.2 cm by 30 cm, ASUS), mounted directly above a graphics tablet (49.3 cm by 32.7 cm, Intuos 4XL; Wacom, Vancouver, WA), as shown in Fig. 1a. Participants made reaching movements while holding a stylus (power grip) embedded within a 3D-printed handle and sliding it along the graphics tablet. The stylus position was recorded at 200 Hz. On each trial, the monitor displayed a yellow start location (6 mm diameter circle), a green circular target (6 mm diameter), and the real-time position of the stylus as indicated by a white circular cursor (3 mm diameter). Both the yellow start location and green target were positioned at the participant’s midline. When seated for testing, participants could not see their arm or hand. The experimental protocol was conducted in the dark to further minimize peripheral vision of the arm. The experimental software was custom written using the Psychtoolbox extension in MATLAB (Brainard, 1997; Pelli, 1997).

### Reaching Task

Participants made quick point-to-point reaches to the straight ahead target located 9 cm from the start position. To initiate each trial, participants moved the stylus so that the white cursor entered the yellow start target. The participant had to maintain their hand in the start position for at least 300 ms (hold time drawn from a uniform distribution, *U* [300, 500]) before the green target appeared 9 centimeters away. Participants were instructed to make quick and accurate point-to-point reaches to the target. If the movement time, defined as the elapsed time from movement onset to movement endpoint, was greater than 500 ms, the target turned red to indicate to the participant that their movement was not quick enough. Movement onset was defined as the first time point that movement velocity was ≥1 cm/s and the hand had traveled 0.5 cm from the center of the start target. Movement end was defined as the first time point following movement onset that the velocity fell below 1 cm/s. The endpoint hand location was taken as the hand position at movement end, with frozen endpoint feedback of the cursor position provided for 500 ms. Following the completion of endpoint feedback, the yellow start target reappeared to prompt the participant to bring the cursor back to “home” to continue the next trial.

### Experimental Schedule

The experiment commenced with a baseline block containing 70 null (unperturbed) trials with veridical feedback. The baseline block was used to familiarize participants with the experimental protocol, ensuring that participants attempted to make quick and accurate point-to-point reaches. Participants were informed that once they started moving, they were to “follow through with their reach to the end” without correcting their movement. We were successful in ensuring participants’ reaction times (282 ms *±* 40 ms; mean of medians *±* SD) and movement times (365 *±* 39) were brisk.

At the end of the baseline block, the experimenter provided the participant with the following instructions: ‘You may notice some changes to your cursor. Regardless of whether you notice these changes or not, continue to aim directly for the target and reach as quickly and accurately as possible in a straight line.’

Trials alternated between null and perturbation trials within each experimental block. On perturbation trials involving a visuomotor rotation, the cursor was rotated away from the hand by 0°, *±*2°, or *±*4° relative to the start position (100 trials / perturbation level).

The participants also experienced trials where there was no visuomotor rotation, but the target jumped by +/- 2°, or +/- 4° mid-reach (hand distance of 4.5 cm). As the focus of the current study is on the results from the visuomotor rotation trials, results from the target jump trials are provided in the Supplement.

Full visual feedback was provided during the reach as well as the return home during all visuomotor rotation trials, with the perturbation left on during both the outbound and inbound portions of the trial. On half of the null trials, randomly selected, visual feedback of the cursor was completely absent during the reach. On these no visual feedback trials, following reach completion, the cursor remained hidden on the return to the start position until the hand was *<*1 cm from the center of the start target. For the other 50% of null trials with visual feedback, participants received full veridical feedback following reach completion and during the return to the start position to begin the next trial. The schedule of perturbation trials was randomized.

Participants completed a total of 18 experimental blocks with 100 trials each, totaling 1800 trials. Each experimental block was separated by a minimum 1-minute break.

### Data Analysis

All data were analyzed using custom-written Python scripts and utilized standard libraries, including NumPy (Harris et al., 2020), pandas (pandas development team, 2020), Matplotlib (Hunter, 2007), and seaborn (Waskom, 2021). Reach angle was defined as the angle between the straight lines connecting the start position to the target and the hand at peak velocity. This measure also quantified IGE, as we only used the straight ahead target at 0°. Structuring the experiment such that each perturbation trial was surrounded by a null trial on either side was critical to our analyses. Single-trial adaptation was quantified as the difference between reach angles on the post- and pre-perturbation trials (adaptation_*t*_ = reach angle_*t*+1_ *−* reach angle_*t−*1_). Operationalizing adaptation this way rather than the more common method of comparing reach angles on post-perturbation to perturbation trials provides an uncontaminated measure of adaptation to internally-generated motor noise. As previously explained (Ranjan, 2022), defining adaptation as reach angle_*t*+1_ *−* reach angle_*t*_ means the outcome measure (adaptation) contains its putative predictor, IGE (reach angle_*n*_), resulting in spurious correlations. We circumvent this potential confound by using the triplet analysis.

For each participant, individual trials were excluded from analysis if the reach angle had an absolute z-score magnitude greater than 3.5. This resulted in *<*1.1% of trials being removed from any individual participant’s data, with 99.5% of all trials in the experiment being included.

We used the following binned analysis method of Ranjan & Smith (see Ranjan, 2022). To estimate sensitivity to EGE, trials were first split based on the level of EGE and further binned into one of five quintiles based on the magnitude of IGE (i.e., 5 levels of EGE x 5 IGE quintiles = 25 bins), separately for each participant. The average magnitude of adaptation within each of these 25 bins was then regressed onto the EGE. For estimating sensitivity to IGE, we regressed the same binned adaptation measures onto IGE. To calculate the *r*^2^ values in Fig. 2, we averaged the 25 bins of adaptation versus EGE (IGE) across participants and regressed the average response onto its respective error signal.

### Statistical analyses

For the analysis of unbinned data, we performed bivariate regression analysis using ordinary least-squares fitting with SciPy’s *linregress* function (Virtanen et al., 2020). After confirming with QQ plots that the data could be modeled as Gaussian distributions, we compared regression coefficients with a paired t-test. We also assessed whether the coefficients were reliably different than zero using a one-sample t-test.

### Modeling Analysis

We separately fit each individual’s dataset with each of the four models using maximum likelihood estimation with SciPy’s *minimize* function (Virtanen et al., 2020). In this procedure, our goal was to find a given model’s parameter set, Θ_model_, that maximized the probability (equivalently, minimized the negative log-likelihood) of the observed trial-by-trial data. We assumed the trials were all independent of each other, and therefore summed the negative log-likelihoods across all trials. For formal model comparisons, we computed and compared Bayesian Information Criterion (BIC) scores (Schwarz, 1978).

To visualize how well each model could capture differential adaptation to EGE-IGE, we generated simulated data, using the actual perturbation schedule, for each participant using each model’s maximum likelihood estimates of parameter values. Fig. 5 shows an example “posterior predictive check” for a representative participant. We note that the patterns observed for this participant were qualitatively similar across our entire sample.

Model recovery analysis with all four models and parameter recovery analysis for PIECE were performed to further validate our model fits (see Supplementary Information; Wilson and Collins, 2019).

Below, we provide details of the models that were implemented.

#### PIECE

In addition to the description of PIECE provided in the main text, we note that the motor prediction, *x*_*u*_, and the proprioceptive cue, *x*_*p*_, are both centered on the true hand location, *x*_*h*_. As we cannot dissociate them, since they are internal to the observer, we combined them, resulting in random variable *x*_combined_ ∼ *N* (*x*_*h*_, *σ*_combined_), where 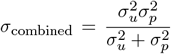. The variance takes this form because *x*_*u*_ and *x*_*p*_ are derived from Gaussian distributions and thus the distribution of *x*_combined_ is the product of these underlying distributions (likelihoods).

In total, there are 3 free parameters in this model: *σ*_combined_, *σ*_*r*_ and *b*, the participant’s intrinsic motor bias.

#### Proprioceptive Recalibration Model (PReMo) (Tsay et al., 2022)

PReMo assumes the computational goal of adaptation is to realign the perceived hand position, *x*_per_, with the target, *T*, which is always equal to zero since we only used the straight ahead target. Computing *x*_per_ on trial *t* involves the following:

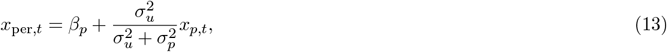

where *β*_*p*_ represents the proprioceptive shift (i.e., how much proprioception gets cross-modally shifted by vision). *σ*_*u*_ represents the uncertainty around the motor prediction, which in PReMo is always centered on the target (in contrast to PIECE, where the prediction is centered on the hand position). *σ*_*p*_ represents proprioceptive uncertainty, and *x*_*p*_ represents the raw proprioceptive input, i.e., the true hand position. The physiological limit of how much proprioception can be shifted is represented by 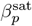, which was set to 5 in order to reduce the number of free parameters:

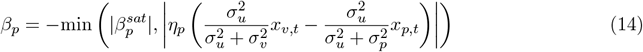

Similar to previous studies that have used randomized perturbations with an average error of zero, we assumed the effects of different perturbations are independent of each other. Therefore, single-trial adaptation (STA) was modeled as resulting only from the current perturbation, with no retention across trials, and is a function of the distance between the perceived hand position and the target. The free parameter *B* represents the learning rate (i.e., error sensitivity):

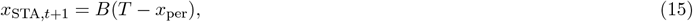

For PReMo and all other models, the actual hand angle on the next trial, *x*_*h*,*t*+1_, is modeled as the sum of single-trial adaptation (learning), the participant’s intrinsic motor bias, *b*, a free parameter, and motor noise, which was taken as a draw, *ϵ*, from a Gaussian distribution, 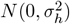 :

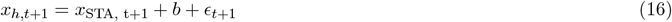

*σ*_*h*_ represents motor variability and is taken directly from baseline reaches. Therefore, there are 6 free parameters in this model: *σ*_*u*_, *σ*_*p*_, *σ*_*v*_, *η, B*, and *b*.

#### Perceptual Error Adaptation Model (Zhang et al., 2024)

For fitting the PEA model to data, we make the same assumptions as the authors of the original work did. Namely, we assume *x*_*u*_ and *x*_*p*_ are, on average, centered on the target position, since we used a randomized perturbation schedule with a mean error of zero. Therefore, instead of serving as separate cues, *x*_*u*_ and *x*_*p*_ were cue combined, resulting in random variable 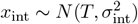, where *T* is the target direction (i.e., zero), and 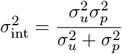 represents the variance of the combined sensory signal from *x*_*u*_ and *x*_*p*_.

Single trial adaptation is a function of the distance between the estimated hand position, 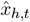 and the target:

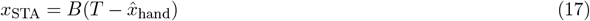

where the estimated hand position is computed by cue combination:

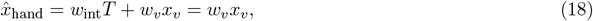

and the weight given to the *i*th cue is given by:

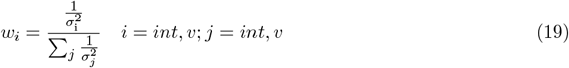

The hand angle on the trial following the perturbation is again given by:

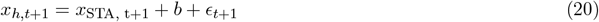

Similar to Zhang and colleagues, we modeled *σ*_*v*_ as a linear function of the cursor distance from the target using the bias (1.179) and slope (0.384) terms from their original work (see Table S1 from Zhang et al., 2024). In total, there were 3 free parameters in this model: *σ*_int_, motor bias, *b*, and the learning rate, *B*.

#### Relevance Estimation Model (REM) (Wei and Körding, 2009)

In the REM model, the observer performs causal inference to compute the probability that the feedback is “relevant” (*p*_rel_), which in this model means the visual error is attributed to the movement made by the observer rather than to the perturbation.

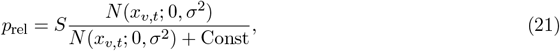

Where *x*_*v*,*t*_ is the visual cue on trial t, S and Const are scaling factors, and *σ* is the standard deviation of the integrated visual and proprioceptive cues, following:

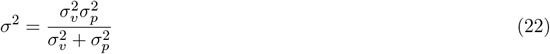

For single trial adaptation:

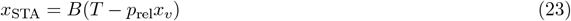

The next motor output is given by:

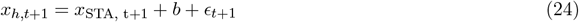

There are 4 free parameters in REM: *σ, S*, Const, and *b*.

## Data and code availability

All data and code for this study are available at the following site: https://github.com/ccmlab-ubc/ige-ege.

## Acknowledgments

The authors thank Maurice Smith, Mike Landy, Todd Hudson, Jonathan Tsay, and members of the Cognition and Action Lab at UC Berkeley for providing helpful feedback on this work. We also thank Lisa Liu for her help with creating Fig. 3.

## Supplementary Information

### Target jump data

We performed the same set of binned and unbinned analyses of participants’ responses to target jumps as we did for visuomotor rotations (see Fig. 2). The results of these analyses are presented in Fig. 7, where each point represents the average across every participant’s mean response for the corresponding EGE/IGE level. Here, EGE refers to the target error and is quantified as the difference between the hand angle at peak velocity and the target angle post-jump. Unlike the visuomotor rotation data (see Fig. 2), there was little-to-no response to either EGE (black dashed line, slope=-0.027 [-0.091, 0.026], *r*^2^=0.094, *p* = 0.136) or IGE (dashed black line, slope=-0.038 [-0.055, 0.014], *r*^2^=0.081, *p* = 0.167). The lack of an adaptive response to the target jumps was further confirmed in our unbinned analysis using a bivariate regression. The sensitivities (i.e., *β*) to EGE (0.027[*−*0.025, 0.091], *t*_15_ = 0.888, *p* = 0.388) and IGE (0.020[*−*0.013, 0.051], *t*_15_ = 1.191, *p* = 0.252) were close to zero. Furthermore, not only were the *β*s for target jumps and IGE near zero, they were not reliably different (mean difference = 0.007, 95% CI for difference scores: [-0.052, 0.066], *t*_15_ = 0.254, *p* = 0.802).

**Figure 7:**
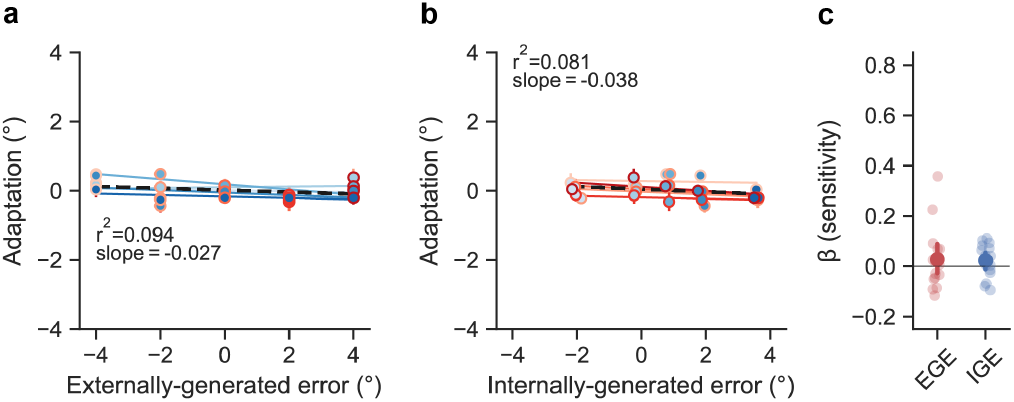
(a) Population-averaged adaptive responses were binned based on the level of EGE (i.e., target error; shade of red) and IGE (shade of blue) and plotted as a function of EGE in a, and as a function of IGE in b (where data are shifted right due to a small counter-clockwise reaching bias of less than 1° across participants). In direct contrast to the visuomotor rotation data, only a minimal amount of the variance in adaptive responses (9.4% here, as compared to 99.1% in the case of rotations) was explained by EGE. (e) Linear regression coefficients from an analysis of unbinned data. Error bars represent bootstrapped 95% CIs and dots represent individual participants.

**Figure 8:**
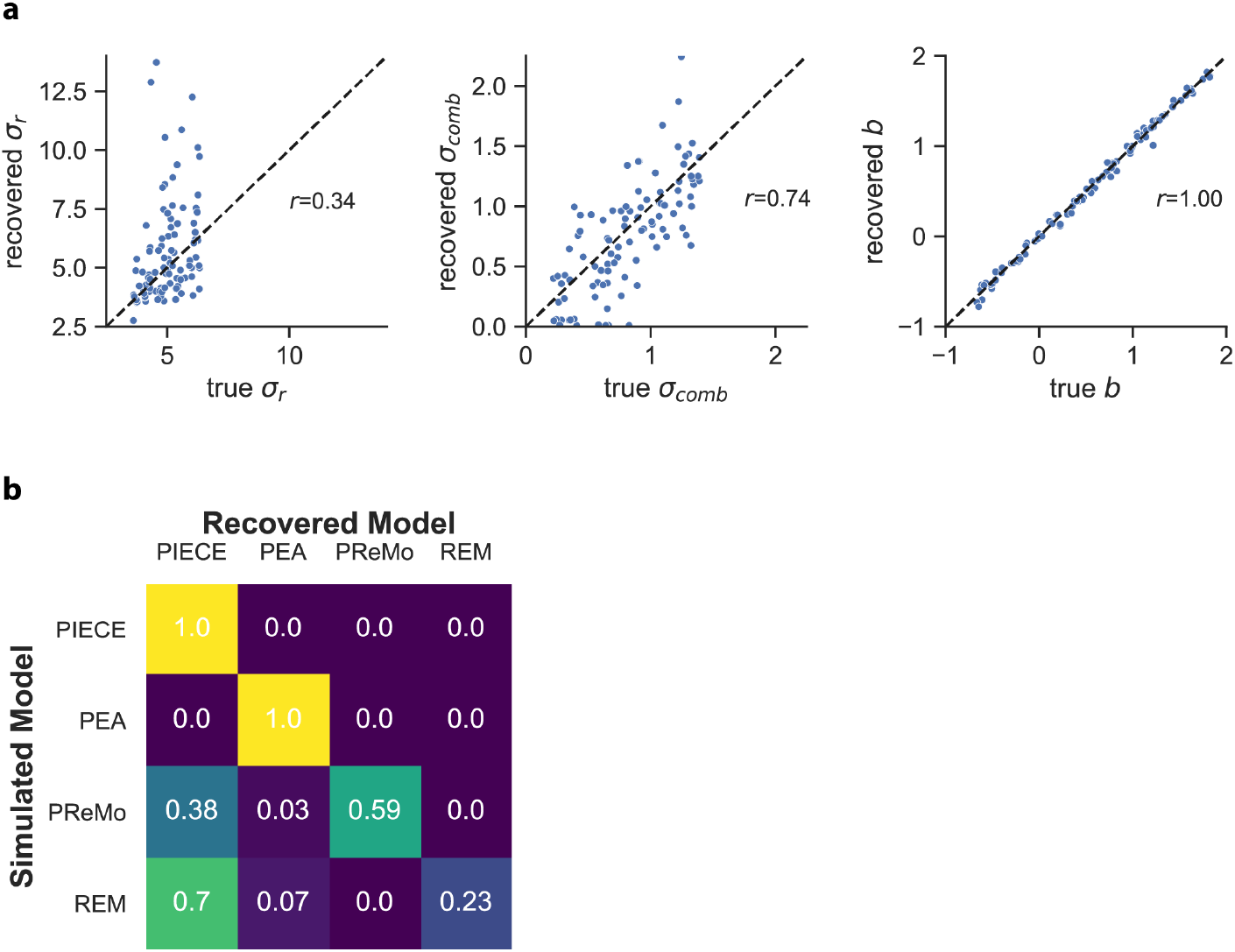
Parameter and model recovery analyses. (a) Plots show the PIECE model parameter values that were recovered (y-axes) from fits to 100 synthetic datasets generated with the PIECE model along with correlation coefficients. The parameter values used to generate the synthetic data are shown on the x-axes. (b) Confusion matrix for model selection. The value of each cell within the matrix indicates *p*(recovered model|simulated model).

Combined, these results from the target jump portion of the experiment indicate that pure target error, where there was no SPE, was not an effective teaching signal for the motor system, consistent with our prior work and the work of others (Oza et al., 2024; Sadaphal et al., 2022; Tsay et al., 2022). Interestingly, these results appear to contrast recent findings that target jumps alone do drive robust implicit adaptation (Ranjan, 2022). However, in our experiment, participants were explicitly instructed to ‘continue to aim directly for the original target’, whether the target jumped or not, thus making the target jump task-irrelevant. In contrast, in the Ranjan & Smith study, quick placement of the hand within the target at the end of each trial was emphasized, thus making the target jump task-relevant. It may be that the implicit adaptation system is sensitive to task-relevant information having to do with target error (i.e., task performance error). This is a potentially interesting dissociation with SPE, where numerous clamped visual feedback studies have shown that participants will robustly adapt to task-irrelevant cursor feedback (e.g., Kim et al., 2018; Morehead et al., 2017; Tsay et al., 2024).

### Parameter Estimates

**Table 1:** MLEs of each model.

| Model | Parameter | Mean [95% CI] |
| --- | --- | --- |
| PIECE | $\sigma_r$ | 5.307 [4.917, 5.697] |
| | $\sigma_{\text{combined}}$ | 0.429 [0.268, 0.589] |
| | $b$ | 0.711 [0.391, 1.032] |
| PReMo | $\sigma_u$ | 2.939 [1.543, 4.334] |
| | $\sigma_v$ | 3.996 [2.461, 5.455] |
| | $\sigma_p$ | 3.958 [2.461, 5.456] |
| | $B$ | 0.575 [0.391, 0.759] |
| | $\eta_p$ | 1 [1, 1] |
| | $b$ | 0.729 [0.374, 1.082] |
| PEA | $\sigma_{\text{int}}$ | 7.028 [5.571, 8.485] |
| | $B$ | 0.297 [0.251, 0.343] |
| | $b$ | 0.858 [0.479, 1.238] |
| REM | $\sigma_{\text{combined}}$ | 6.538 [4.824, 8.251] |
| | $S$ | 0.944 [0.357, 1.531] |
| | $Const$ | 2.755 [0.904, 4.606] |
| | $b$ | 0.848 [0.476, 1.219] |

### Parameter and Model Recovery

To validate the parameter values obtained by fitting PIECE to our behavioral data and our ability to distinguish between each of the four models, we performed parameter recovery and model recovery analyses (Wilson and Collins, 2019). We generated 100 synthetic datasets with each of the four models used in this study, totaling 400 datasets in all. The synthetic data from each model were generated using parameter values drawn from uniform distributions bounded by the minimum and maximum parameter values obtained from fits to each individual participant’s behavioral data (i.e., the range of MLEs of each model). We then fit all 400 synthetic datasets with each model using maximum likelihood estimation and performed objective model selection for each dataset.

As shown in Fig. a, the recovered parameters for *σ*_comb_ and *b* from the PIECE model were highly correlated with the parameter values used to generate the synthetic data. In the case of *σ*_r_, the correlation was not as strong. Future experiments that test a wider range of perturbation sizes will result in more accurate recovery of this parameter, as it represents the observer’s prior uncertainty about the perturbation size, which in the current experiment only spanned a very narrow range.

More importantly, our model recovery analysis showed that we can accurately identify whether data were generated by the PIECE or PEA model, and to a lesser extent with PReMo. Based on BIC scores, PIECE best fit the synthetic data generated with REM more frequently than REM did. This points to PIECE being able to accommodate a wide range of adaptation behaviors, and is also due to PIECE being a slightly more parsimonious model than REM (3 free parameters for PIECE versus 4 for REM). Notably, the results of our model recovery analysis were qualitatively similar when using Akaike Information Criterion scores, with the only difference being that the diagonal values in our confusion matrix were larger for PReMo and REM using AIC scores. This points to the robustness of our modeling results, especially with regard to PIECE being clearly identifiable.

